# Nonlinearities and Timescales in Temporal Interference Stimulation

**DOI:** 10.1101/2022.02.04.479138

**Authors:** Tom Plovie, Ruben Schoeters, Thomas Tarnaud, Luc Martens, Wout Joseph, Emmeric Tanghe

**Author notes:** Correspondence: Tom Plovie.

## Abstract

In temporal interference (TI) stimulation, neuronal cells react to two interfering sinusoidal electric fields with a slightly different frequency. It was previously seen that for the same input intensity, the neurons do not react to a purely sinusoidal signal. This study aims to get a better understanding of the mechanism underlying TI neuromodulation, which is largely unknown. To this end, single-compartment models are used to simulate computationally the response of neurons to the sinusoidal and TI waveform. This study compares different neuron models to get insight into which models are able to reproduce the experimental observations. It was found that integrate- and-fire models do not entirely reflect the experimental behavior while the Hodgkin-Huxley and Frankenhaeuser-Huxley model do reflect this behavior. Changing the characteristics of the ion gates in the Frankenhaeuser-Huxley model alters the response to both the sinusoidal and TI signal, possibly reducing the firing threshold of the sinusoidal input below that of the TI input. The model results show that TI stimulation is not qualitatively impacted by nonlinearities in the current-voltage relation. In contrast, ion channels have a significant impact on the neuronal response. This paper makes advances both in terms of biophysical insight into the neuron as well as the insight in computational modelling of TI stimulation.

## 1 INTRODUCTION

Nowadays, many applications for brain stimulation exist. One of these applications is deep brain stimulation (DBS) that was mainly developed to treat movement disorders, such as Parkinson’s Disease (PD). However, it has an important disadvantage: it is invasive and can cause complications (e.g. bleeding in the brain, stroke, infection) (Aum and Tierney, 2018). Because of this, a new technique is being developed named temporal interference (TI) stimulation. With this technique, two sinusoidal electric fields, oscillating at a high but slightly different frequency, are applied to the brain by two cathode-anode pairs. At the place where they temporally interfere, an amplitude-modulated signal is created. This is illustrated in Figure 1. The frequency at which the envelope repeats is called the beat frequency. Grossman et al. (2017, 2018) has found from experiments on mice that cortical and hippocampal neurons respond to this interference pattern but not to the constituting sinusoids separately. Therefore, TI stimulation is a non-invasive alternative to the conventional, invasive DBS techniques. Furthermore, instead of using electrodes to create an amplitude modulated electric field inside the brain, it is also possible to use transcranial magnetic stimulation (TMS) coils, creating a magnetic field and inducing an electric field. It is possible to achieve a TI waveform by using the same technique of applying multiple frequencies to the brain (Sorkhabi et al., 2020; Xin et al., 2021). Although promising, there are still some limitations to overcome before TI stimulation can be used for humans and can replace conventional DBS. A first limitation is that the technique has only been extensively tested on rodents, while very few studies are performed with experiments on humans. One study reports TI stimulation of the human hippocampus with improved episodic memory accuracy as a result (Violante et al., 2022). Another study used TI stimulation to show an increase in activity in the striatum and in its functional brain network. This resulted in a better performance in a motor learning task, with a prominent improvement for healthy older subjects Wessel et al. (2022). Furthermore, Vassiliadis et al. (2022) showed the importance of the striatum in reinforcement learning processes via the use of TI stimulation. A second limitation is that the mechanism in the neuron that allows this type of stimulation is unknown. It is thus very hard to predict how different neuron types will react to the stimulation and thus how exactly the human brain responds to TI stimulation.

**Figure 1.**
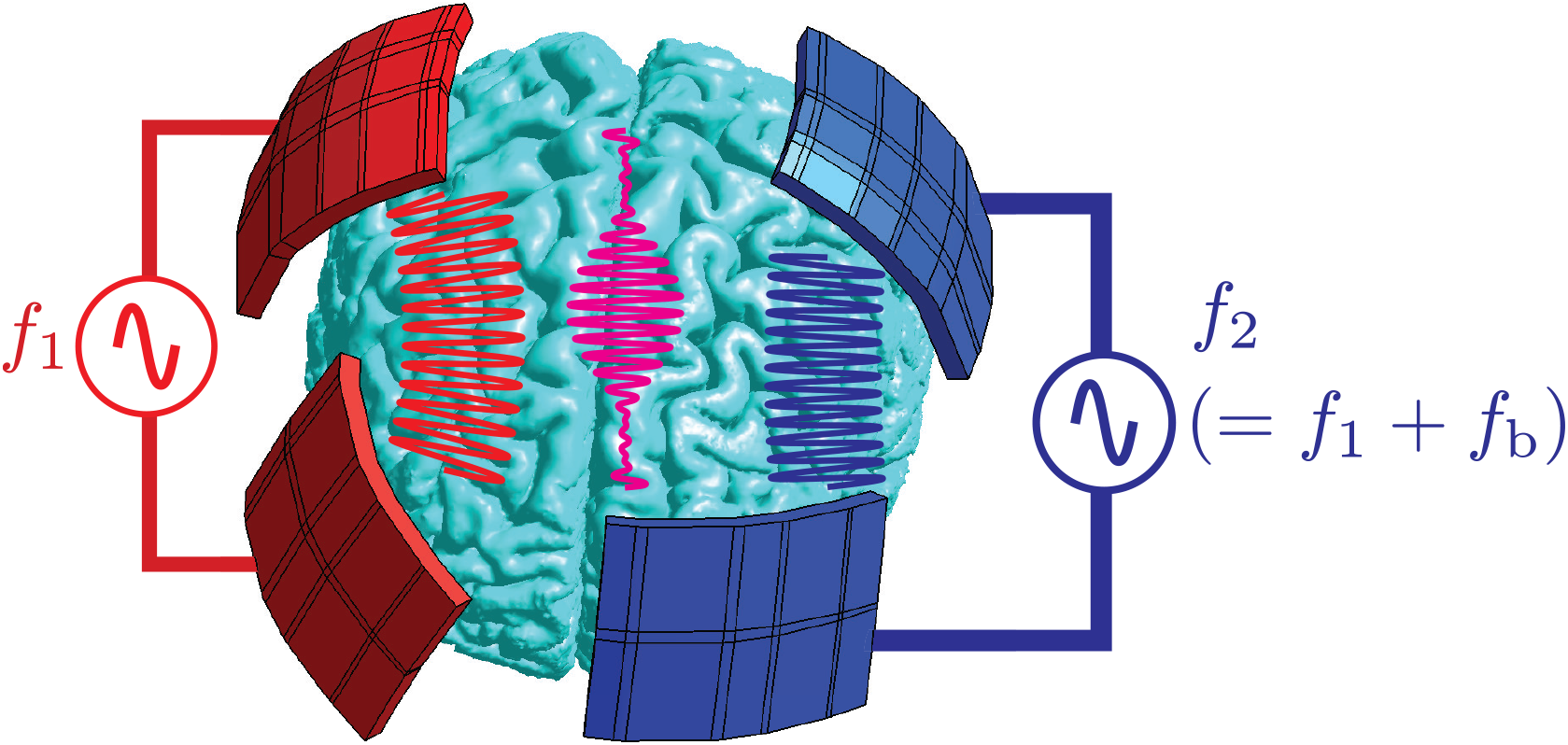
Schematic illustration of the concept of TI-DBS in the MIDA brain(Iacono et al., 2015). The red and blue electric field oscillate at a different frequency. At the place where they temporally interfere, the pink amplitude modulated signal is created.

Other experiments in mice have shown that it is possible to cause eye movements at the beat frequency of the TI signal (Song et al., 2021), confirming the importance of the beat frequency. It should be noted that TI stimulation is not only used in the brain. It was shown in rats that TI stimulation can activate spinal motor neurons in the neck to restore breathing after opioid overdose. Again it was shown that a single sine waveform is not able to cause stimulation (Sunshine et al., 2021). Furthermore, TI has been studied for retinal stimulation (Su et al., 2021).

To get more insight into the electric field distribution of TI stimulation in the human brain, finite element method (FEM) simulations have been performed. In these studies, TI stimulation has been compared to a conventional technique like transcranial alternating current stimulation (tACS). This showed that when the same electric field strength is achieved inside the focus, TI induces a smaller maximal electric field outside this focus than tACS (von Conta et al., 2021). In contrast to the traditional DBS technique, it is easier to steer the stimulation site with TI stimulation. This can be done by changing the electrode positions or by adjusting the relative amplitudes of the current at the different electrode pairs (Rampersad et al., 2019; von Conta et al., 2021; Gomez-Tames et al., 2021). With these degrees of freedom, it possible to find the optimal electrode positions for targeting focality. This optimal electrode positioning is very person-specific. With the same electrode positioning, high variations are possible in the simulated electric fields for different people (von Conta et al., 2021). Huang and Parra (2019) argue that it is possible to achieve sufficiently high electric fields in deep brain regions using tACS. To achieve this, the electrode positioning becomes more important because it makes use of the cerebrospinal fluid as a conductor for the electric field. It is thus even more challenging to obtain a patient-specific electrode position. They do mention that the focality is still worse with this technique than with TI stimulation (Huang and Parra, 2019). The focality in TI stimulation is in the centimeter range (Huang et al., 2020). Lee et al. (2022) proposed epidural TI stimulation, where the electrodes are placed under the skull, in order to avoid stimulation in cortical regions. They validated their FEM simulations with phantom experiments. Furthermore, the feasibility of TI stimulation in humans is investigated by Acerbo et al. (2022) with the use of mice models and human cadavers in combination with simulations for the optimal electrode positioning.

It was found in silico and in vivo that increasing the carrier frequency of the TI signal increases the threshold of the input needed to cause stimulation. Changing the beat frequency did not seem to have an influence on the threshold according to Grossman et al. (2017) and Gomez-Tames et al. (2021). On the other hand, Mirzakhalili et al. (2020) and Howell and McIntyre (2021) do mention the importance of the beat frequency for the activation threshold. Furthermore, Plovie et al. (2022) showed that for a reduced Hodgkin-Huxley model, the excitation threshold changes with both carrier and beat frequency. To model TI stimulation, it was found that integrate-and-fire (IF)-neurons are not sufficient to reproduce the experimental behavior on single neuron level (Cao et al., 2020). However, when IF-neurons are placed in a network and GABA_b_ postsynaptic inhibition is included, the computational results were in line with experimental observations of TI stimulation (Esmaeilpour et al., 2021). Further on, for point neurons, it is possible to have neurons exhibiting TI stimulation when using the FitzHugh-Nagumo model or the Hodgkin-Huxley model. It was also seen that not all mammalian cells exhibit TI (Cao and Grover, 2018; Cao et al., 2020). Howell and McIntyre (2021) argues that because of the large dimensions of a human head, TI stimulation seems unfeasible. However, it is suggested that subthreshold TI stimulation can be used to synchronise the firing of neurons to the TI signal. Next to the immediate effect on the neuron, TI stimulation could also, like other electrical stimulation methods, impact the plasticity and the interconnectivity of neurons (Howell and McIntyre, 2021). Up to now, it was said that high-frequency signals do not cause stimulation. This is not entirely true: there are actually three ways for a neuron to respond to a high-frequency signal. There can be either one action potential (AP) at the signal onset, continuous firing or depolarization block. It may be impossible to stimulate the deep regions of the brain without also causing a conduction block in other parts of the brain (Mirzakhalili et al., 2020). This conduction block occurs at higher input intensities and could lead to an upper limit for the use of TI stimulation (Grossman et al., 2017). The conduction block was also seen in Wang et al. (2022), with morphologically-realistic neuron models at amplitudes that were an order of magnitude larger than the amplitude for activation. This again shows the difficulty with off-target regions during TI stimulation (Wang et al., 2022).

Regarding the underlying mechanism at the neuron level that enables this stimulation type, one of the earliest hypotheses was that TI stimulation can be explained in terms of the intrinsic low-pass filter property of a neuron together with a nonlinear component. For example, in the Hodgkin-Huxley framework, the cell membrane is modeled by a parallel RC circuit, thus implementing a low-pass filter when driven by a current source Hodgkin and Huxley (1952). Here, a first nonlinearity is introduced in the time-dependent resistor, through the ion channel dynamics. Indeed, the channel opening and closing rate functions depend nonlinearly on the membrane potential. Furthermore, subsequent work has generalized the Ohmic current-voltage relation in the Hodgkin-Huxley model to account for the nonlinear distribution of electrolyte concentration along the ion channel Frankenhaeuser and Huxley (1964). The resulating Goldman-Hodgkin-Katz current-voltage relation introduces a second nonlinearity to the equation set. Because of the low-pass filter property, the neuron cannot react to the high-frequency oscillations and therefore responds to the low-frequency envelope of the TI signal (Grossman et al., 2017, 2018). It is clear that the low-pass filter property cannot be the single explanation because with only the properties of a low pass filter, the envelope could not be extracted by the neuron and it would not respond to the beat frequency. This is because the TI waveform, as a sum of two high-frequency sines, does not contain spectral content at the beat frequency. As a result, a pure low-pass filter will not give spectral components at the beat frequency. Alternatively, by first rectifying the signal and then low-pass filtering, which comes down to an envelope detection of the TI signal, the beat frequency is extracted. In fact, a rectification process is needed in order to have a response to the TI-stimulus. However, the neuronal membrane cannot be a perfect demodulator because high-frequency oscillations can still be found in the membrane potential response (Mirzakhalili et al., 2020; Cao and Grover, 2018).

The first finding of this paper is the demonstration that the nonlinearity of the Goldman-Hodgkin-Katz (GHK) equation in the Frankenhaeuser-Huxley model is not strictly required to demonstrate the effect of TI stimulation found in Grossman et al. (2018). Consequently, our model results indicate that envelope detection can be realized at the membrane level through the nonlinearity intrinsic to ion channel gating. From the comparison of biologically inspired models (Frankenhaeuser-Huxley and Hodgkin-Huxley) and different IF models, a second finding is the required aspects in a computational neuron model to show the effect of TI stimulation. The third novel aspect is a thorough investigation of the different influences of both the time constants and the steady-state parameters of the gates to obtain a biophysical explanation of the neuronal response to TI stimulation. It was already mentioned by Mirzakhalili et al. (2020) and Cao et al. (2020) that the time constants influence the excitation threshold. In this study, this is investigated in more detail.

## 2 METHODS

### 2.1 Neuron models

In this study, single-compartment models are used. This means that the morphology of the neuron is not taken into account. This can be justified as the idea is to investigate the influence of nonlinearities on the neuronal response to sinusoidal signals and TI signals. The longitudinal direction of the neuron does not add any nonlinearities and is thus not thought to have a big influence on the underlying mechanism of TI stimulation. The five existing models that are used are the Hodgkin-Huxley (HH) (Hodgkin and Huxley, 1952), Frankenhaeuser-Huxley (FH) (Frankenhaeuser and Huxley, 1964), leaky integrate-and-fire model (LIF), exponential integrate-and-fire (EIF) and the adaptive and exponential integrate-and-fire (AdEx) model. The first two are described by the following equations:

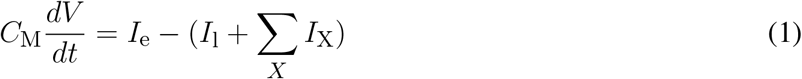

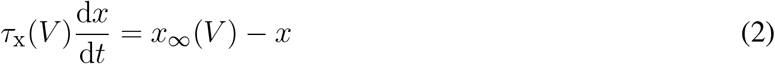

where *C*_M_ is the membrane capacitance, *V* is the membrane potential, *I*_e_ is the input current, *I*_l_ is the leak current and *I*_X_ are the currents linked to voltage-dependent gates, where *X* can be Na, K or p (sodium, potassium and a non-specific current, respectively). The values for all of the constants are taken from Hodgkin and Huxley (1952) and Frankenhaeuser and Huxley (1964). In (2), *x* can be *m, h, n* or *p*. These are the open probabilities of the respective gates. *p* is only used in the FH-model. *τ_x_*(*V*) and *x*_∞_(*V*) are the time constant and the steady-state value characterising the gates. These are again retrieved from Hodgkin and Huxley (1952) and Frankenhaeuser and Huxley (1964). For TI, *I_e_* is of the form 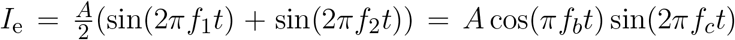 is the beat frequency and *f_c_* is the carrier frequency. The ion currents of the FH-model and HH-model are of the form of (3) and (4), respectively.

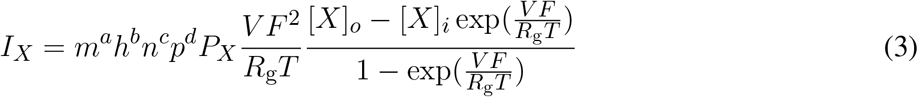

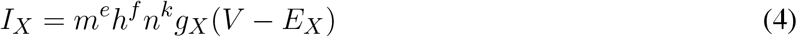

The part of (3) after the gates (*m*, *h*, *n* and *p*) is known as the Goldman-Hodgkin-Katz (GHK) flux equation. *P_X_* is the ion-specific permeability, *F* is the Faraday constant, *R*_g_ is the ideal gas constant and *T* is the temperature. The exponents *a, b, c, d, e, f* and *k* are the number of gates of the respective channel and depend on both the model and the ion type. [*X*]_o_ and [*X*]_*i*_ are the extracellular and intracellular concentrations of the ion X, respectively. The HH- and FH-model are made for the squid giant axon and the myelinated nerve fibre of Xenopus laevis, respectively. However, mammalian neurons function in a similar way. As this study focuses on the qualitative behavior of neurons, these models are well suited for the aims of this study.

The LIF-model is described by:

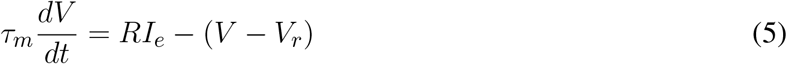

and the EIF-model is described by:

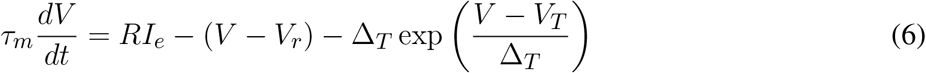

where *τ_m_* is the membrane time constant, *R* is the membrane resistance, *V_r_* is the resting potential, Δ_*T*_ is the slope factor that determines the sharpness of the threshold and *V_T_* is the model intrinsic spike threshold. When the threshold *V*_thresh_ is reached, the potential resets to *V*_reset_. Note that the LIF-model is the EIF-model in the limit of Δ_*T*_ → 0. The AdEx-model is very similar but has an additional differential equation. It is described by:

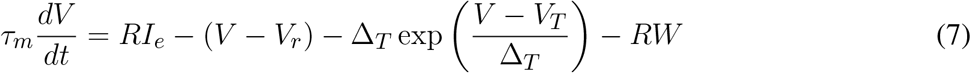

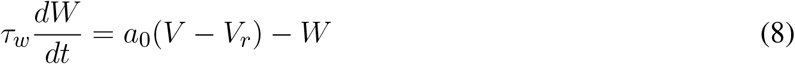

where the additional parameter *a*_0_ determines the level of subthreshold adaptation. Again when a threshold *V*_thresh_ is reached, the potential is reset to *V*_reset_ while *W* is changed into *W* + *b*_0_ (Brette and Gerstner, 2005). The parameters used for the presented results of the IF-models are given in Table 1 unless stated otherwise. The IF-models are mainly added in order to investigate which characteristics are needed in neuron models to make the neuron spike at lower inputs for a TI signal than for a sinusoidal signal.

**Table 1.** Parameters for the IF-models

|  |  |
| --- | --- |
| $V_r$ | $-70$ mV |
| $V_{\text{reset}}$ | $-70$ mV |
| $V_{\text{thresh}}$ | $-40$ mV |
| $R$ | $0.5 \cdot 10^6$ kΩ |
| $\tau_m$ | $5$ ms |
| $\Delta_T$ | $2$ mV |
| $V_T$ | $-50$ mV |
| $\tau_w$ | $100$ ms |
| $a_0$ | $2 \cdot 10^{-9}$ mS |
| $b_0$ | $8 \cdot 10^{-6}$ μA |

### 2.2 Neuron model extensions

To investigate the influence of the nonlinearity of the GHK-equation, two new models are constructed from the HH- and FH-model. A partly linearized version of the FH-model (FH1) is obtained by replacing the GHK-factor of (3) by its first-order Taylor expansion for *I_K_*, *I_Na_* and *I_p_*. In (9), this is shown more specifically by the transition from left to right.

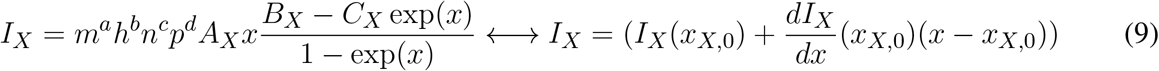

where 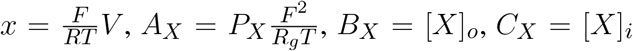 and 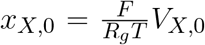. In this transition, the gates remain the same as in the original models. The ion currents are thus still nonlinear because of the gate equations, only the part representing the GHK-equation is linearized for FH1. For the HH-model, the opposite is done. It is assumed that the HH-model is the first-order Taylor expansion of a more general model. This more general model is then of the form of (3) and is called the gHH-model. In (9), this corresponds to the transition from right to left. In order to obtain these new models, the right expansion points ((*V*_*Na*,0_, *V*_*K*,0_) and (*V*_*Na*,0_, *V*_*K*,0_, *V*_*p*,0_)) for this Taylor series should be found for the gHH and FH1 model, respectively. Two methods are followed to realise this, resulting in four derived models: gHHv1, FH1v1 (method 1) and gHHv2, FH1v2 (method 2).

The first method consists of calculating the reversal potentials for the ion currents (i.e. the Nernst potentials) and taking these as the expansion points. In the second method, a sweep is done over several combinations of possible Taylor expansion points. For each set, several combinations of duration and intensity of a rectangular input signal are tested. The values for the input amplitudes are ten uniformly distributed values between 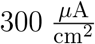 and 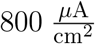 for the FH-model and between 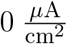 and 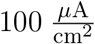 for the HH-model. For the stimulation duration, ten uniformly distributed values between 0.05 ms and 2.00 ms are chosen for both models. For every combination, the correlation between the membrane response of the original and the new model is calculated. The correlations of the different duration-intensity combinations are averaged. The (*V*_*Na*,0_, *V*_*K*,0_)-combination with the highest average correlation is chosen for the new model. For the FH1 models, it is assumed that *V*_*p*,0_ = *V*_*Na*,0_.

### 2.3 Gates

To investigate the influence of the time constants, a small adaptation is made to (2). A parameter *k_x_* is introduced to scale the time constant of the gates so that

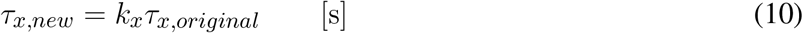

Lastly the steady-state values of the gates are investigated in more detail. The equations for the steady-state values are replaced by a logistic function like (11), while the original time constant equations are maintained.

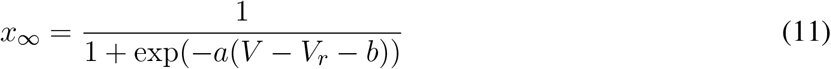

The influence of the parameters *a* and *b* on the TI-effects can then be investigated. The goal of this approach is to get insight into how the behavior changes with changing parameters and not to approximate the real behavior of the neuron globally. *V_r_* is the rest potential and is equal to −70 mV.

### 2.4 behavior analysis

The analysis of the results was done based on the firing rate (FR) and the new energy metric defined below in (13). To obtain the firing rate, first the action potentials (APs) need to be determined. For the IF-models, an AP is counted every time the threshold *V*_thresh_ is exceeded. For the HH-like models, an AP was detected based on the m-gate. The m-gate, that is responsible for the *Na*^+^-influx that initiates the AP, needs to cross a threshold value of 0.75 for an AP to be counted. This definition is preferred because the membrane potential oscillates with the carrier frequency, while the m-gate is more robust. An additional rule is set that the m-gate needs to drop below 0.10 before a new AP can be counted, in order to avoid miscounts when the m-gate oscillates around 0.75. To cancel out the influence of transient behaviors caused by the onset of the input current, only a certain time window is considered to determine the FR. This time window is from 100 ms after the stimulation onset until 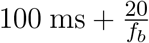 after the stimulation onset, where *f_b_* is the beat frequency. In this way, 20 periods of the envelope are included. Then the FR is calculated with:

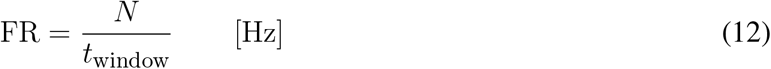

where *N* is the number of APs counted and *t*_window_ is the duration of the stimulation in the investigated time window, being 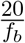.

A disadvantage of the FR metrix is that it depends on the arbitrary lower and upper excitation threshold values of 0.10 and 0.75. Because it is not ideal that the results depend on the choice of these values, a metric based on the energy of the signal is introduced for extra analysis. This metric pTI (percentage of TI) is defined as:

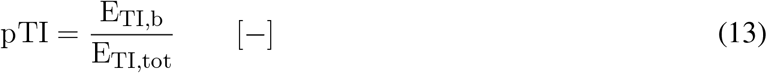

where *E*_TI,b_ is the sum of the energy at the beat frequency and the first harmonic of the beat frequency in the frequency spectrum of the membrane potential in response to the TI signal. *E*_TI,tot_ is the total energy in the membrane response to the TI signal (i.e. the sum of the magnitudes at all frequencies). The total energy is restricted to energies at frequencies higher than 0 Hz in the frequency spectrum. The DC-component is thus not taken into account, to ensure that pTI is independent of the resting potential.

### 2.5 Software

The programming, analysis and simulations were performed in MATLAB R2021b. The ode45, ode15s and ode4 solvers of Matlab were used to calculate the neuronal response numerically (Shampine and Reichelt, 1997). For example, the fixed step ode4-solver was needed for transforming the signals to the frequency spectrum. The variable step variable order solvers, ode45 and ode15s were used to speed up the simulations when no frequency information was needed. The principle of the ode45 and ode15s solvers is explained in Ashino et al. (2000).

## 3 RESULTS

A summary of the most important results is given in Tables 2 and 3. Table 2 shows for which of the discussed models it is possible to get a TI zone. The TI zone is defined as the range of input intensities for which the TI signal causes regular firing and the sine input with the same carrier frequency does not cause regular firing. Regular firing is defined as the neuron having more than one spike during the considered time window explained in section 2.4.

**Table 2.** Summary of the existence of a TI zone for the discussed models. EIF includes both the EIF1 and EIF2 model.

|  | yes | no |
| --- | --- | --- |
| LIF | x |  |
| EIF |  | x |
| AdEx |  | x |
| HH | x |  |
| FH | x |  |
| gHH | x |  |
| FH1 | x |  |

**Table 3.** Summary of the gate parameter values, for which there is a TI zone in the FH model with a carrier frequency of 3 kHz and a beat frequency of 100 Hz.

| | $a$ [1/mV] | $b$ [mV] | $k$ [-] |
| --- | --- | --- | --- |
| $m$ | whole range | $>27.8$ | $<1.58$ |
| $h$ | whole range | $<18.7$ | whole range |
| $n$ | whole range | whole range | whole range |

### 3.1 IF-models

Figure 2 shows the threshold for regular firing, induced by a TI waveform (solid lines) and by a single sinusoid at the same carrier frequency (dashed lines). The input current is divided by an area of 1 cm^2^ to get a measure in 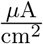. In the case of the LIF model, the TI signal causes APs at a lower threshold than the sinusoidal signal. The LIF model thus shows a TI zone. However, the width of this zone is on average only 0.0646% of the threshold of the TI signal. For both EIF models, the single sinusoid causes regular firing at a lower threshold than the TI signal. The TI threshold is 1.13 and 1.18 times higher than the sine threshold for the EIF1 and EIF2 model, respectively.

**Figure 2.**
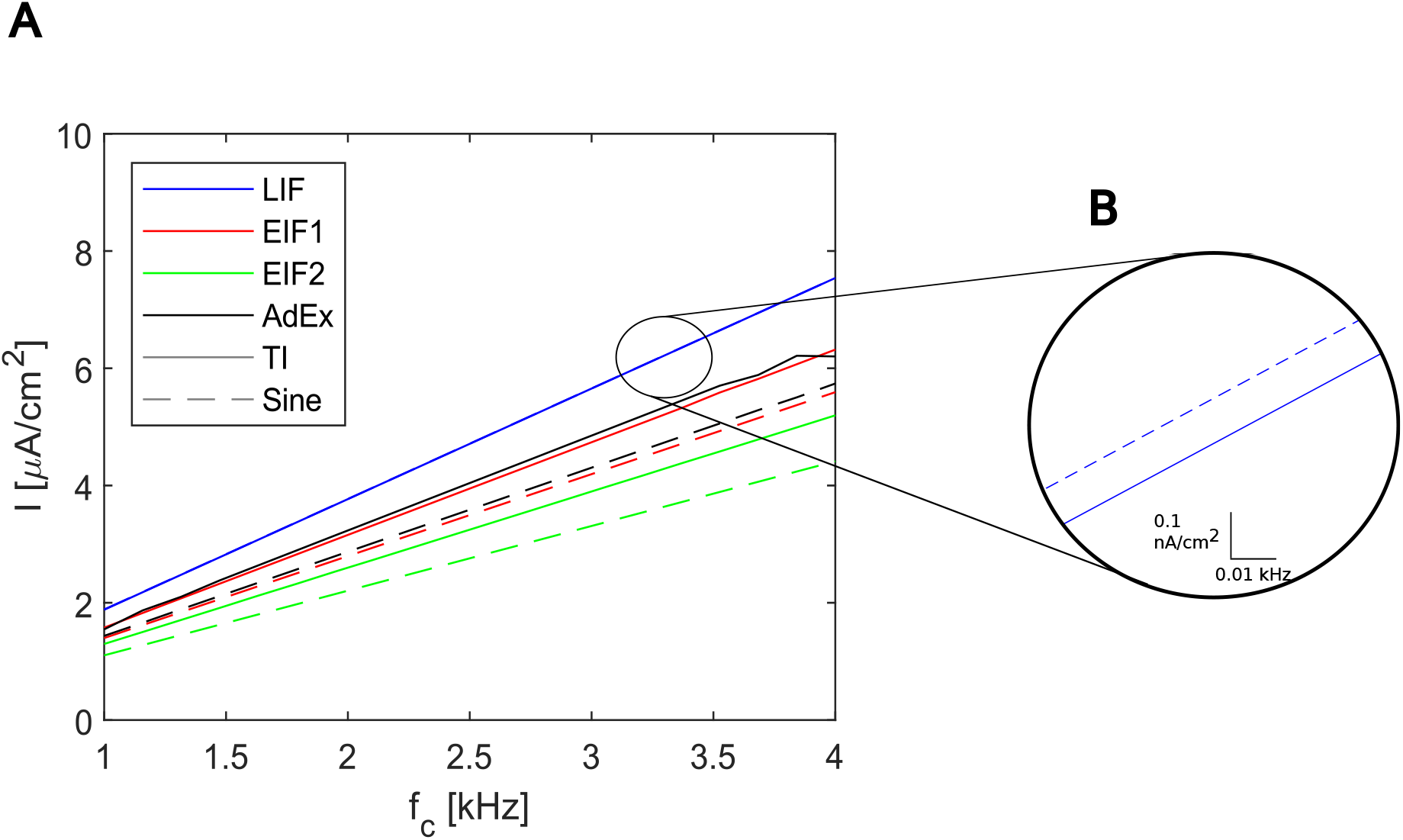
Threshold lines for regular firing with a TI input (solid lines) and a sinusoidal signal (dashed lines) with carrier frequency *f_c_* and a beat frequency of 100 Hz. The thresholds are shown for the leaky integrate-and-fire (LIF), exponential integrate-and-fire with Δ_*T*_ = 2 mV (EIF1), exponential integrate-and-fire with Δ_*T*_ = 15 mV (EIF2) and adaptive and exponential integrate-and-fire (AdEx) neuron. **A**: overview of all IF models. **B**: detail of the LIF model.

For the AdEx-model, the TI signal again causes firing at higher intensities (approximately 1.12 times higher for a TI signal than for a sinusoidal signal). For all models and all signals, it is clear that for a higher carrier frequency, a higher input is needed to achieve stimulation. None of the IF models showed a depolarization block over the range of 1 kHz – 4 kHz.

### 3.2 Effect of nonlinearity in current equations

Two methods are used to determine the Taylor expansion points for the gHH and FH1 models, as described in section 2.2. For the first method, the expansion points are chosen equal to the Nernst reversal potentials of the ion currents. For the HH-model, this results in *V*_*Na*,0_ = 45 mV and *V*_*K*,0_ = –82 mV. For the FH-model this results in *V*_*Na*,0_ = 53.53 mV, *V*_*K*,0_ = –97.74 mV and *V*_*p*,0_ = 53.53 mV.

For the second method, the average correlations are calculated as described in section 2.1. The maximum for the FH1v2-model can be found at (*V*_*Na*,0_ = –30.69 mV, *V*_*K*,0_ = –37.24 mV) and is equal to 0.994. The average correlation for the FH1v1-model is 0.924. For the gHHv2-model, the maximum is at (*V*_*Na*,0_ = 0.34 mV, *V*_*K*,0_ = –78.6 mV) and is equal to 0.971. The average correlation for the gHH1v1-model is 0.785. Figure S1 in the supplementary information shows the average correlation as function of the Taylor expansion points for the gHHv2 and FH1v2 models. Some examples of voltage traces are shown in Figure S2.

#### 3.2.1 FH vs FH1

Figures 3A-C show the energy plot of the FH-model and the difference with the energy of the FH1-models (pTI_FH1_ – pTI_FH_) as a comparison. In Figure 3A it can be seen that for a higher *f_c_* value, there is a larger range of input intensities for which the neuron reacts at the beat frequency of the TI-input. Also for a higher *f_c_*, a higher input is needed to achieve spiking. Moreover, it can be seen that the behavior of the FH1-models is very similar to the original FH-model (Figures 3B-C). Here, FH1v2 is a closer match than FH1v1 to the original FH-model. The only difference is that the FH1-models get high energies at the beat frequency at higher input intensities.

**Figure 3.**
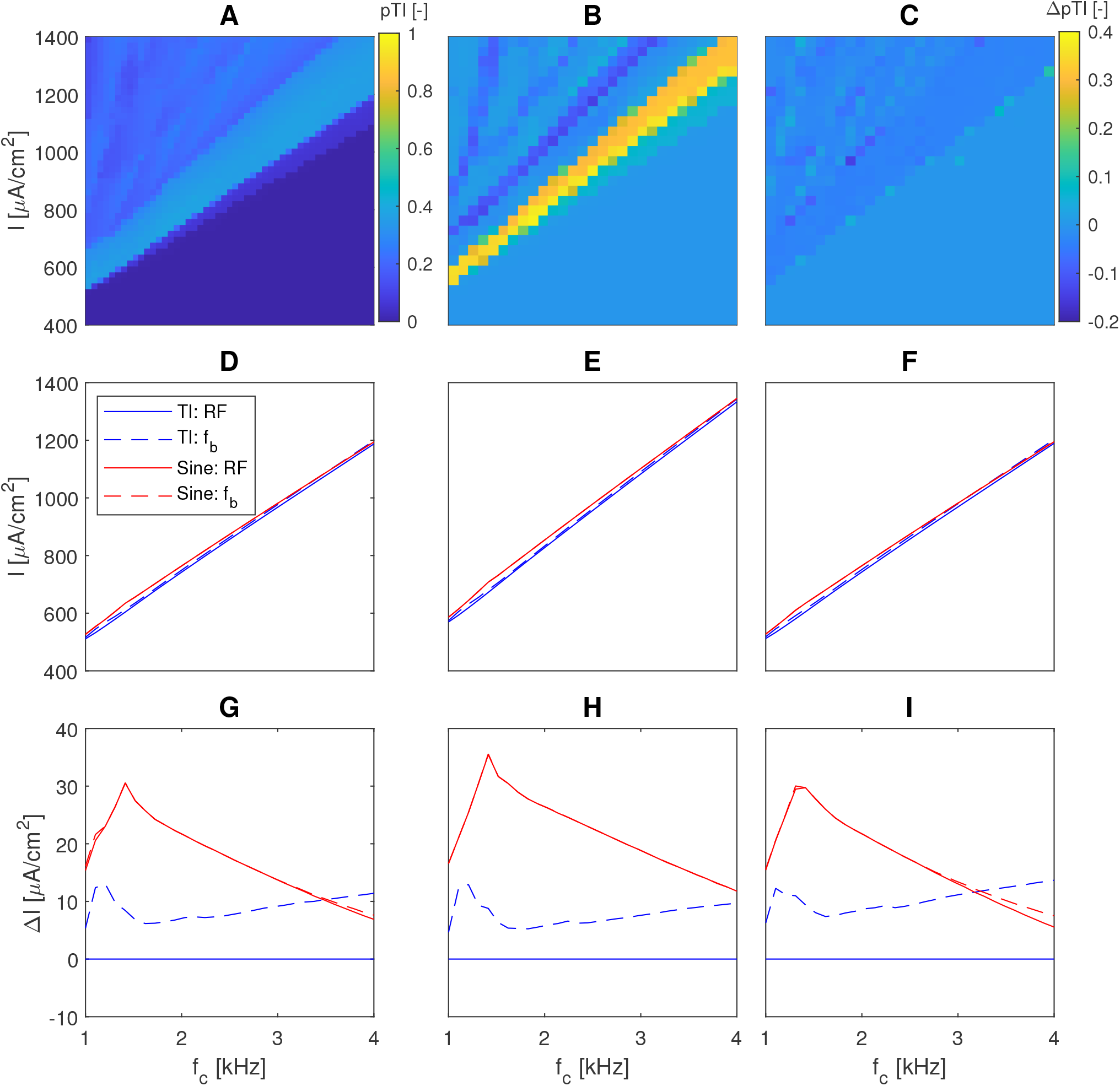
Comparison of the FH- and FH1-model with a beat frequency *f_b_* = 100 Hz. **A**: 2D plot of the energy metric for the FH model **B-C**: Difference between the energy metric of the FH1v1 and FH1v2 model with respect to the FH model, respectively. **D-F**: Absolute threshold lines for TI and sinusoidal stimulation to obtain regular firing (RF) and firing at the beat frequency (*f_b_*) in the FH, FH1v1 and FH1v2 models, respectively. **G**-**I**: Relative threshold lines with respect to the threshold line of regular firing (RF) of the TI-input in the FH, FH1v1 and FH1v2 models, respectively.

The plots of the threshold lines can be used to compare the models as well. These results are shown in Figures 3D-I. From the threshold lines, the TI zone can be deduced. Again it can be seen that the models are very alike with some minor differences (maximal relative difference between thresholds of 15.03 % and 3.49 % for FH1v1 and FH1v2 with respect to FH, respectively). Comparing the two solid lines, the FH model and FH1 models have a TI zone in the whole range of the carrier frequency from 1 kHz to 4 kHz. The maximal width of the TI zone is 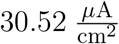, 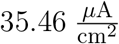 and 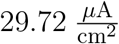 for the FH, FH1v1 and FH1v2 model, respectively. The FH1v2-model again shows the best match to the FH-model over the whole *f_c_*-range. Considering the fact that these TI zones are very small, an example is shown in the supplementary information with a larger TI zone (Figure S3). In this case, the *a* and *b* parameters of the logistic function are changed (cf. (11)). The influence of these parameters are discussed in sections 3.3.2 and 4.4.2.

Moreover, it can be seen that higher intensities are needed to obtain firing at the beat frequency, compared to regular firing. For the FH model the additional intensity needed is 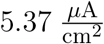 up to 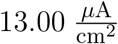 in the range of 1 kHz to 4 kHz. Finally it is interesting to consider the FR for the sine at regular firing. For *f_c_* > 1.2 kHz, these FRs are always above 80 Hz.

#### 3.2.2 HH vs gHH

The results for the energy metric and the firing thresholds for the HH and gHH models are shown in Figure 4. From the energy plots it seems that the models again react very similar with only some minor differences due to the approximation. The firing thresholds however suggest that there are big differences for a sinusoidal input. For the sinusoidal input in the HH model, the neuron never spikes above a carrier frequency equal to 1.74 kHz, while for the gHH-model spiking is seen over the whole *f_c_*-range. It is thus possible that the GHK-nonlinearity assures more robustness for quickly oscillating signals. Regarding the TI zone, it is visible that in the whole range in which the pure sine is able to cause APs, the TI signal has a lower threshold. For the gHHv1 model, there is only a TI zone for *f_c_ >* 1.65 kHz. The gHHv2 model again shows a TI zone over the whole *f_c_*-range between 1 kHz and 3 kHz. The maximal width of the TI zone is 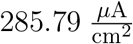, 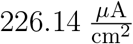 and 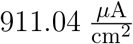 for the HH, gHHv1 and gHHv2 model, respectively. Interesting is the threshold for firing at the beat frequency. This threshold of the sinusoidal input is lower than that of the TI input for certain parameter choices. This applies for the HH model for *f_c_* < 1.4 kHz, in the whole tested *f_c_* range for the gHHv1 model and for *f_c_* < 2.63 kHz for the gHHv2 model.

**Figure 4.**
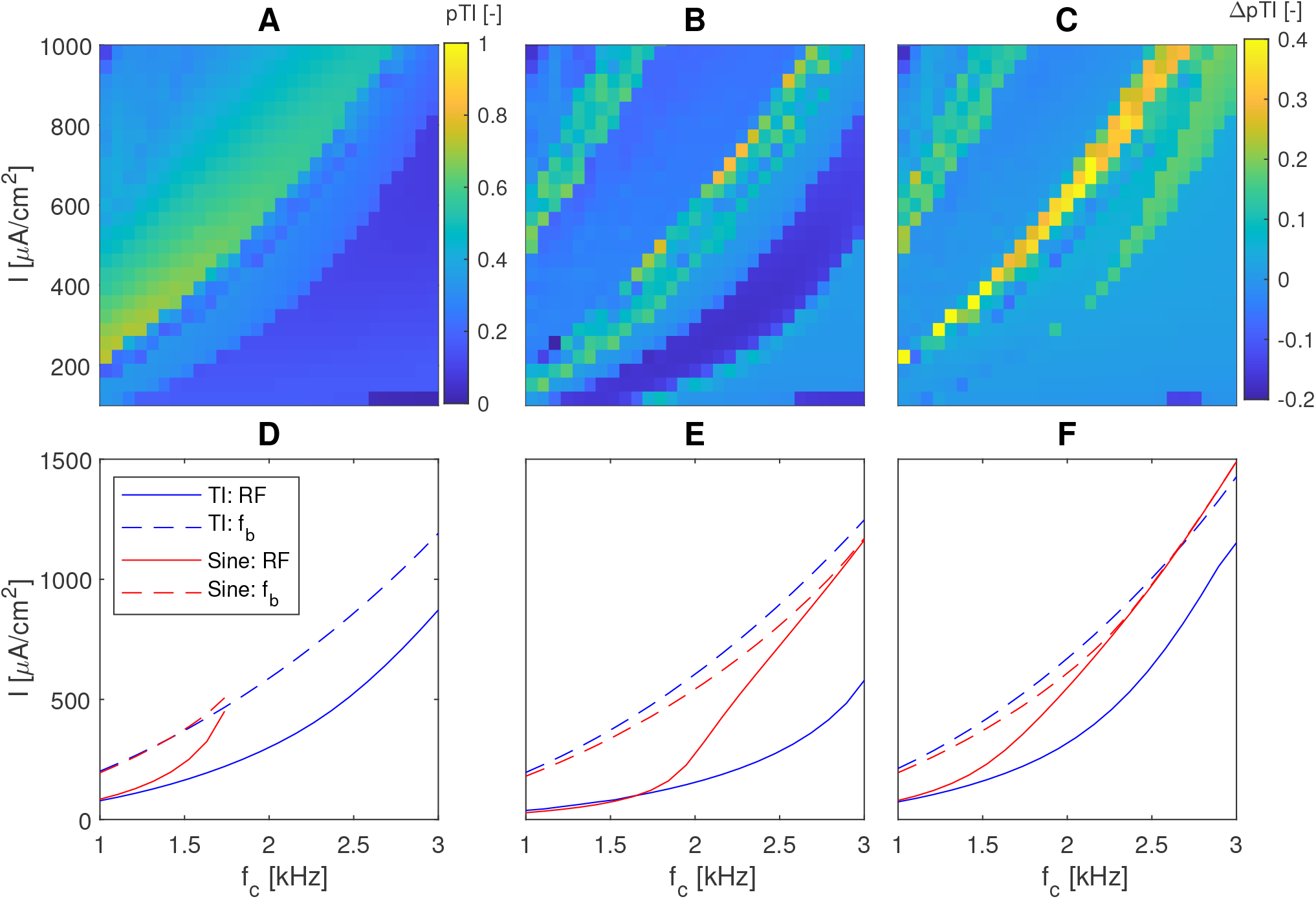
Comparison of the HH- and gHH-model with a beat frequency *f_b_* = 100 Hz. **A**: 2D plot of the energy metric for the HH model. **B-C**: Difference between the energy metric of the gHH1v1 and gHH1v2 model with respect to the HH model, respectively. **D-F**: Threshold lines for regular firing (RF) and firing at the beat frequency (*f*_b_) of the HH, gHHv1 and gHHv2 models, respectively.

### 3.3 Gates

#### 3.3.1 Time constants

Figure 5 shows the influence on the firing thresholds for the different *k*-parameters. For the m-gate (Figure 5A), a logarithmic trend is followed. The thresholds increase with increasing *k_m_*-values until it reaches a plateau. Moreover, the blue and red lines cross around *k_m_* = 1.57, meaning the TI zone disappears for higher *k_m_*-values.

**Figure 5.**
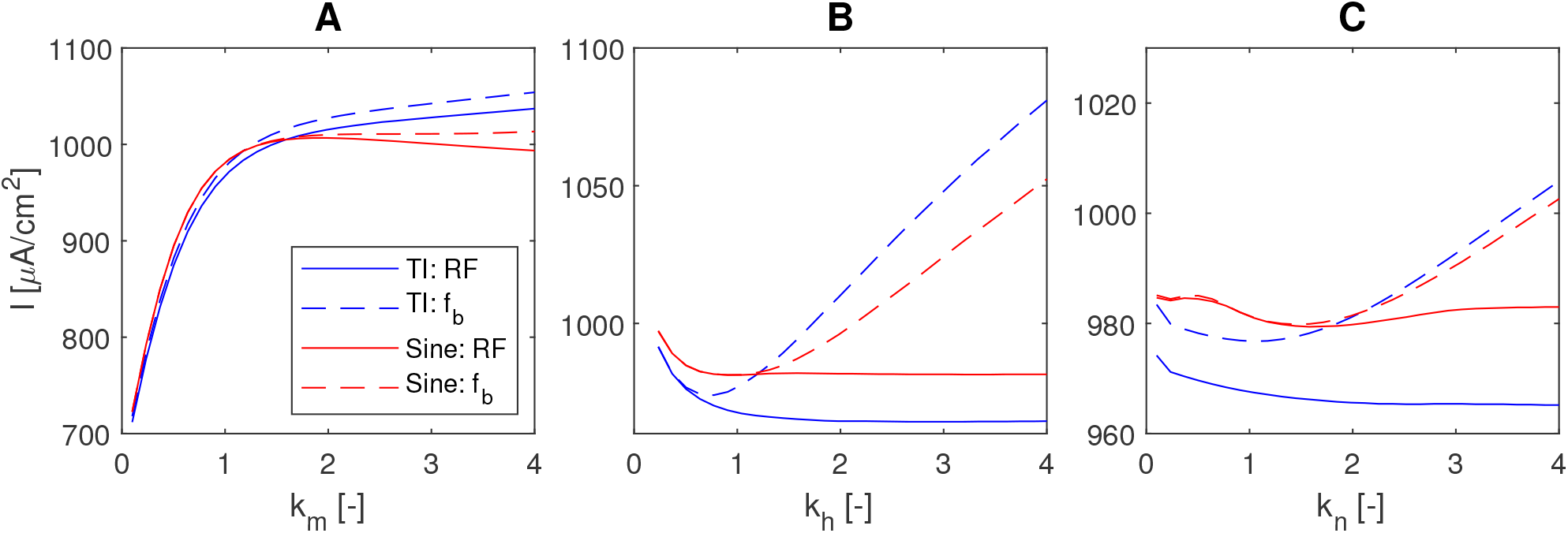
Comparison of the threshold lines for regular firing (RF) and firing at the beat frequency *f*_b_ = 100 Hz for different *k_x_*-values for the FH model (*f*_c_ = 3 kHz). **A**: m-gate. **B**: h-gate. **C**: n-gate.

For very low values of *k_h_* (< 0.7), the input intensity needed to achieve spiking increases rapidly for both the TI and sine signal for decreasing *k_h_* values. When *k_h_* < 0.1, it is not possible anymore to create APs. For *k_h_* > 0.7, the threshold lines increase for firing at the beat frequency but stay approximately constant for regular firing.

Similar results are found for the n-gate: for very low *k_n_*-values (*k_n_* < 1), all input thresholds increase, while for high *k_n_*-values the thresholds stay more or less constant except for firing at the beat frequency.

Furthermore, it can be seen that the TI signal is able to cause regular firing at lower input intensities than needed for firing at the beat frequency. For low k-values, the solid and dashed blue line in Figure 5 are almost equal. In contrast, a gap of 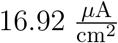, 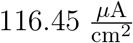 and 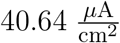 arises for *k_m_* = 4, *k_h_* = 4 and *k_n_* = 4, respectively.

Finally, it can again be seen that for certain *k* values the threshold for firing at the beat frequency is higher for the TI signal than for the sinusoidal signal. This is the case for *k_m_* > 1.18, *k_h_* > 1.22 and *k_n_* > 2.09.

#### 3.3.2 Steady-state values

The effect of changing the slope of the m-gate’s membrane potential dependency is shown in Figure 6A. A higher a-value shifts the TI zone to higher input currents. The influence of the b-value (i.e. shifting the logistic function along the x-axis) of the m-gate is similar (Figure 6B). A small increase in the b-value makes the threshold lines shift to higher input currents. Moreover, a plateau is observed for *b* < 27 mV where the neuron fires spontaneously.

**Figure 6.**
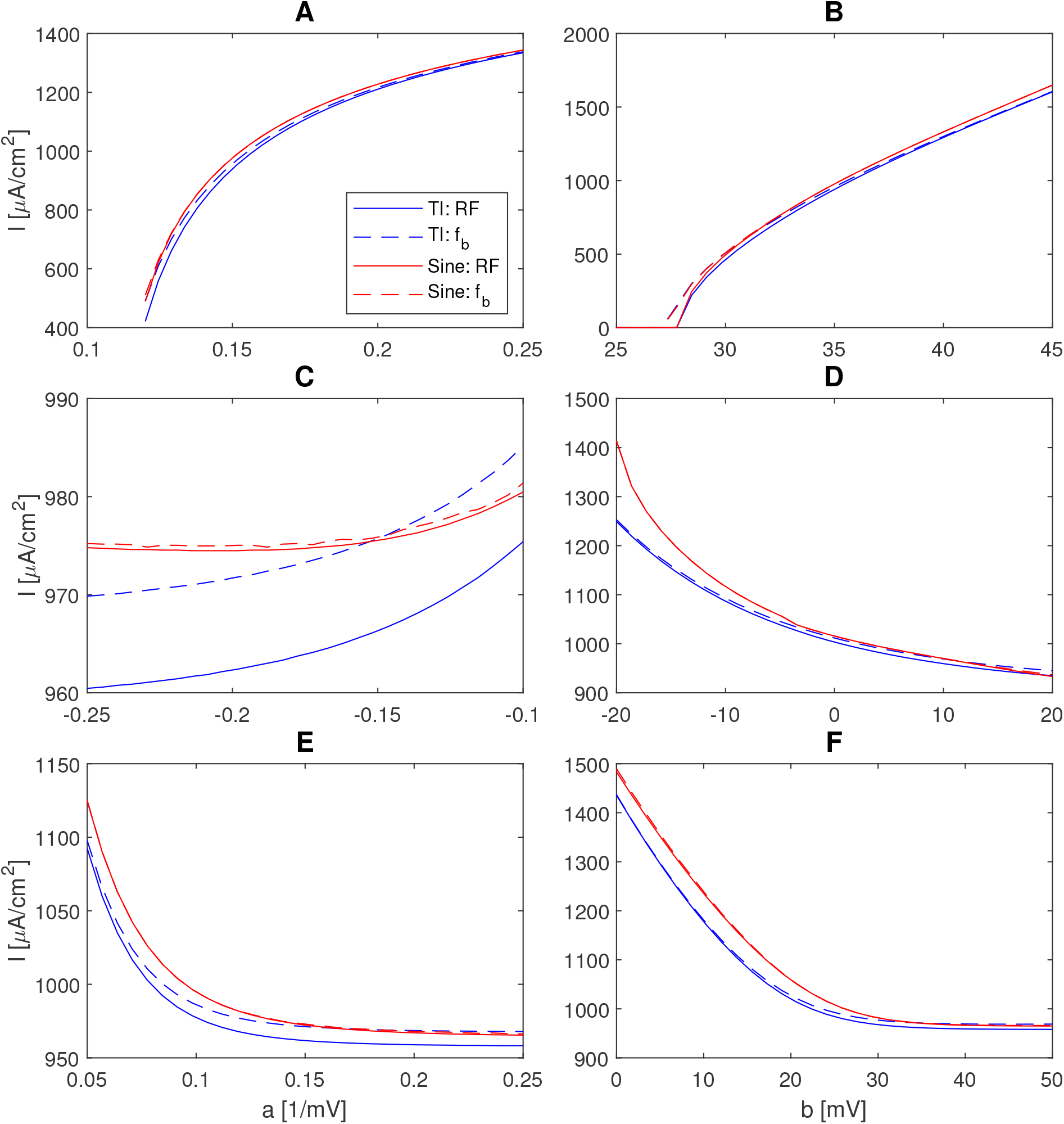
Results for the threshold lines for the FH model with *x*_∞_ as a logistic function with a varying *a*-parameter and constant *b*-parameter (left) and a varying *b*-parameter and constant *a*-parameter (right) with *f_c_* = 3 kHz and *f_b_* = 100 Hz. **A-B**: m-gate. **C-D**: h-gate. **E-F**: n-gate.

For the h-gate, a similar but opposite effect is seen. An increase in b-value shifts the thresholds, and thus the TI zone, to lower input currents (Figure 6D). It has to be noted that this effect is less pronounced. While for the m-gate, a slight change in *b*-value causes a big shift along the y-axis, for the h-gate a bigger change in *b*-value is needed to accomplish the same shift. Increasing the *a*-parameter of the h-gate increases the threshold for spiking activity. In contrast to the *m*-gate, it increases in a convex way. An interesting observation is that the threshold for the TI input increases faster than for the sine-input with an intersection of the two dashed lines at 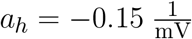. Also for the b-parameter, an intersection of the dashed blue and red line can be seen at *b_h_* = 11.48 mV.

In case of the n-gate, increasing the *a*-value shifts the threshold lines to lower input currents. The TI zone spans over the whole *a_n_* and *b_n_* ranges, but there are intersections of the dashed lines for 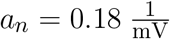 and *b_n_* = 34.48 mV (Figure 6E-F).

The dependency of the existence of a TI zone on the steady state values and time constants is summarized in Table 3.

## 4 DISCUSSION

### 4.1 IF models

Besides the LIF model, for all investigated IF models it was seen that no TI-behavior is present (Figure 2). Although the LIF model does not contain any nonlinearities, it is still possible to get a TI zone, despite being small. The reason for this is that the TI signal contains both the *f*_1_ and *f*_2_ frequencies, while the sine only contains the *f_c_* frequency. Because *f*_1_ < *f_c_*, the threshold is slightly lower for the TI signal than for the pure sinusoid. This is further explained with the analytical result in the supplementary information (section S2). From the transition from LIF to EIF it is clear that adding a nonlinearity makes the TI zone disappear. The transition from EIF to AdEx did not show major differences. However, in Esmaeilpour et al. (2021), AdEx neurons did show correspondence with the experiments of Grossman et al. (2017). In that study, they used a network of neurons rather than a single neuron. They also state that the effect was probably seen because they added the slow GABA_b_ channels to the network, while isolated AdEx neurons are unlikely to exhibit the TI behavior.

### 4.2 FH and HH model

Two conclusions can already be drawn from the FH and HH models. First, the threshold for the neuron to exhibit regular firing or to fire at the beat frequency is not equal. Second, the FR for a sinusoidal input is around 100 Hz in most cases. The reason for the first conclusion will be discussed more in depth in the next sections. The second conclusion points out the importance of TI stimulation. When applying a TI input, it much easier to control the FR of the neurons than for a sinusoidal input, for which the FR changes much faster with the input intensity. For pure sines, lower FRs can then only be reached when the current is applied in a pulsed way. If indeed the difference between the excitation thresholds of a TI signal and a sinusoidal signal is very small, the controllability of the FR with the TI signal can still be an advantage for TI stimulation as in Howell and McIntyre (2021) and Wessel et al. (2022), where it is proposed to use a subthreshold TI signal.

### 4.3 Importance of the GHK nonlinearity

From the comparison between the FH and FH1 models it became clear that linearising the GHK-equation of the ion currents has a minor effect on the neuronal response. However, the TI zone does not disappear as seen with the IF-models. Instead, the TI zone behaves as an approximation of the more complex model without losing its most important characteristics. Simply adding a nonlinearity in the differential equation of the membrane potential is thus not sufficient to fully change the behavior of the neuron model. The shift of the pTI-metric to higher input intensities is explained by the observation that the *Na*^+^-current is lower in magnitude for the FH1-models, so a higher input is needed to achieve firing. This is supported by the fact that the FH1v2-model is a better approximation in which the magnitude of the *Na*^+^-current is higher, especially at lower membrane potentials.

The comparison of the HH and gHH models shows larger differences. For example, it is less straightforward to determine which linearised model better approximates the behavior of the HH-model. However, with both models it is possible to achieve a TI zone. From this it can be concluded that the GHK nonlinearity is not the main reason for a neuron model to show a TI zone.

### 4.4 Gates

The effect of the different characteristics of the gates will be explained in this section, but the mechanism of TI can be best explained with Figure 7. Figure 7 shows that a TI signal can elicit an AP at every other beat, while the sinusoidal signal with the same carrier frequency can only elicit an AP at the stimulation onset. An explanation for this behavior can be found by examining the gates. For the response to a sinusoidal signal, all the gates reach a state of small-amplitude oscillations about a quasi-equilibrium value. However, the values around which they oscillate are not favorable for neuronal excitation (a higher h-value and a lower *n*-value would be better). Comparing this to the TI signal (Figure 7A), it can be seen that because this input signal drops to lower values for a longer period, the n- and h-gates are able to return to values that are more favorable to create an AP. This is further illustrated by the observation that no AP is induced by the second beat in Figure 7A, because the h-gate value is still too low. However, at the subsequent beat, further opening of the h-gate results in an AP.

**Figure 7.**
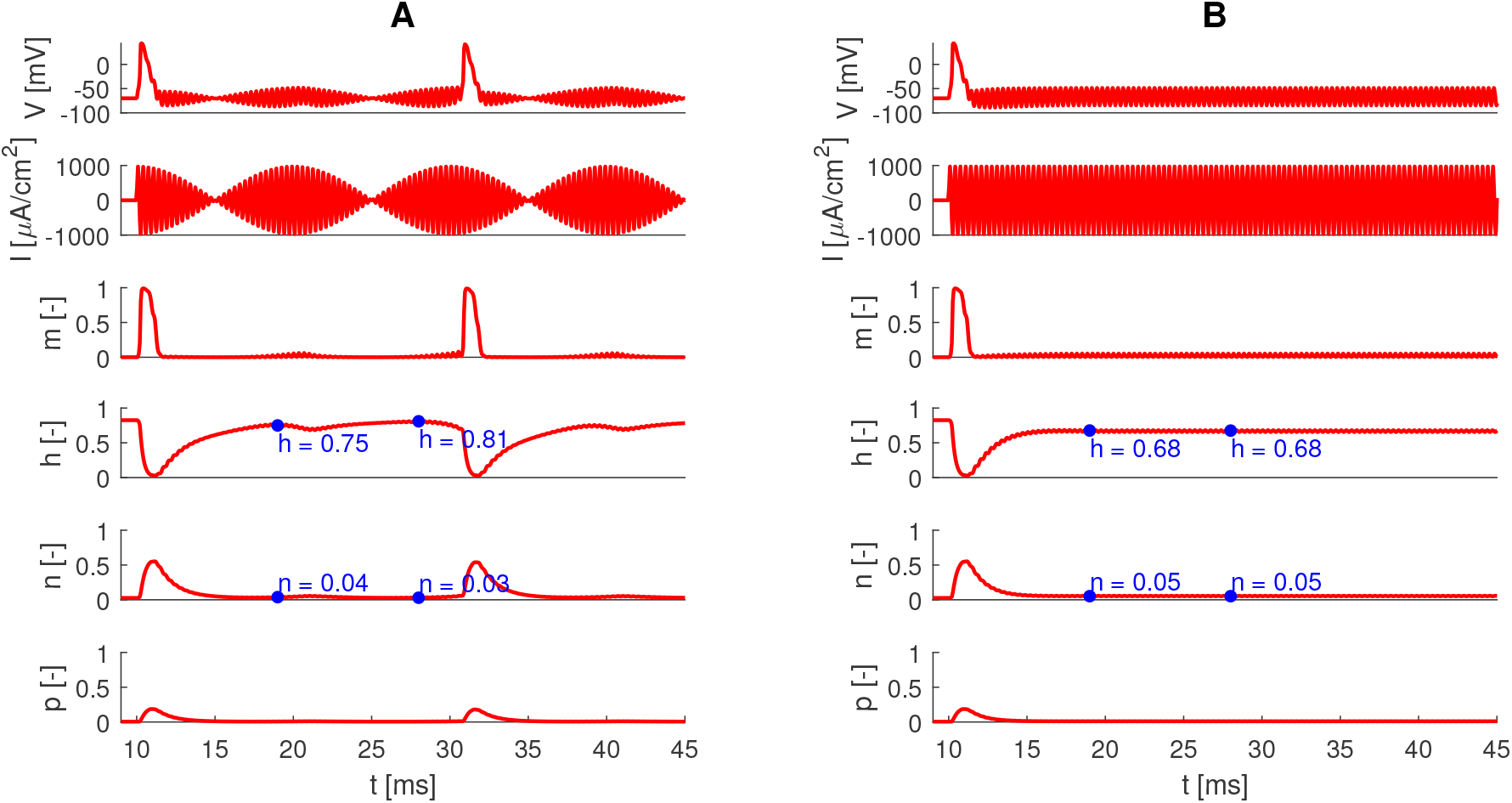
Illustration of the different responses of an FH-neuron to a TI signal (*f_b_* = 100 Hz, *f_c_* = 3 kHz and 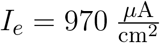) and a sinusoidal signal (*f* = 3 kHz and 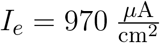). **A**: TI-input. **B**: sinusoidal input.

#### 4.4.1 Time constants

The results in Figure 5A show that the m-gate time constant has the most influence on the behavior of the neuron. For lower time constants, the neuron starts to fire at lower amplitudes of the input current. This can be explained by the fact that the fast reaction of the m-gate is responsible for the initiation of an AP. If the m-gate would react infinitely fast, an AP would be created at the moment that a certain threshold of the membrane potential is reached. When the m-gate reacts with a certain lag, a higher membrane potential should be reached to create an AP, as the m-gate needs some time to get to a value that is high enough to push the membrane potential into an AP. However, if the potential starts decreasing again shortly after its maximum is reached and the m-gate starts decreasing as well, no AP will be created. Because of this, the firing thresholds increase proportional to the m-gate time constant (cf. Figure 5A). Above *k_m_* = 1, the threshold for regular firing of the sine decreases slightly. This is due to an interplay of the closing and opening of the m-gate over multiple periods of the sine.

For the h-gate it was seen that both very low and very high time constants increase the threshold current for firing at the beat frequency (Figure 5B). In the case of a very low time constant, this is because the h-gate closes too fast (Figure 8A). In contrast, high time constants result in reduced h-gate opening after a first action potential, resulting in higher excitation thresholds (Figure 8B). This also explains why from a certain point, increasing *k_h_* only has an influence on the threshold for firing at beat frequency of the TI signal (dashed blue line). The slower the h-gate reacts, the more beats are skipped and the higher the input current has to be to force the neuron to respond to every beat. This will not affect the regular firing threshold of the TI signal because the effect is only on the time between spikes. When the h-gate has reopened, the rapid response of the m-gate will always be able to initiate a new AP, only the FR will be lower. For sine-input, the same reasoning can be made: waiting long enough for the h-gate to reopen will always lead to a new AP, only with a lower FR.

**Figure 8.**
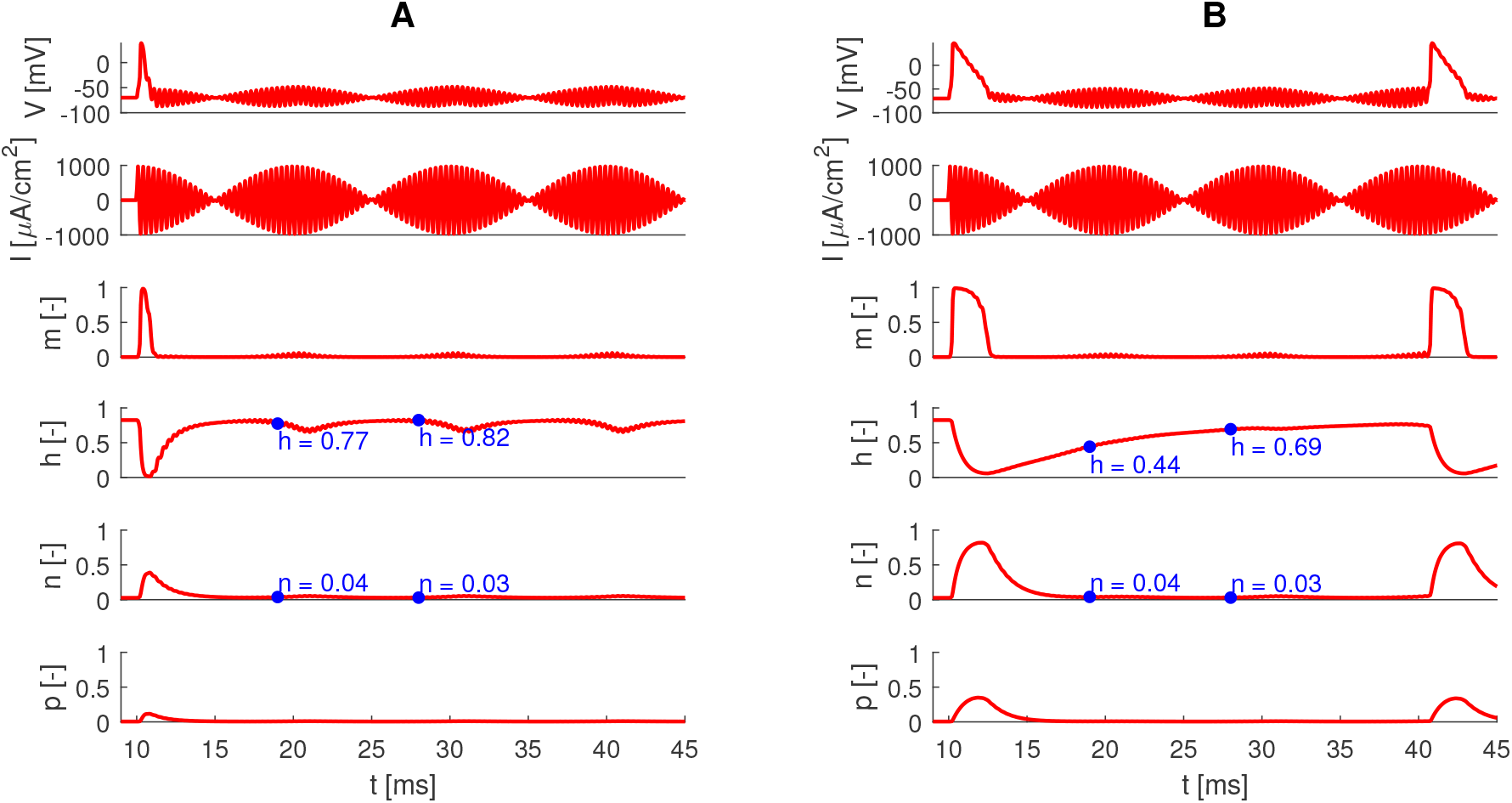
Illustration of the different responses of an FH-neuron to a TI signal (*f_b_* = 100 Hz, *f_c_* = 3 kHz and 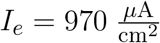) with different *k_h_*-values. **A**: *k_h_* = 0.5. **B**: *k_h_* = 3.

The reasoning for the n-gate is very similar to that of the h-gate. Opening of the n-gates causes an outflow of *K*^+^-ions resulting in a decrease of the membrane potential. Both gates can thus counteract the formation of an AP with the only difference that the effect of the h-gate is present in the form of a multiplication with the m-gate, while the effect of the n-gate is present in the addition of the *K*^+^-current. Therefore, a fast-reacting n-gate (low *k_n_*) makes it harder to create an AP and thus a higher input current is needed to achieve spiking (cf. Figure 5C). For a slower n-gate (high *k_n_*), the *K*^+^-outflow will still be too high to initiate a new AP at the next beats. The reasoning why the threshold for firing at the beat frequency of the TI signal changes most with increasing *k_n_* is the same as with increasing *k_h_*.

The threshold for firing at the beat frequency is lower for a sine input than for a TI input for *k*_m_ > 1.18, *k*_h_ > 1.22 or *k*_n_ > 2.09. This is because increasing the input intensity above the threshold for spiking for a sine will increase the FR in a continuous way. In contrast, increasing the intensity above the threshold for a TI signal will not immediately increase the FR. The FR will only increase when more beats can cause an AP. Therefore, the FR will make a discrete jump when increasing the input intensity. Moreover, at the theshold, the neuron is expected to fire around the trough of the envelope. When the input is just too low, it is possible that some beats do not elicit an AP, because the states of the gates were not right in the short time frame around the top of the beat. In this way, it is possible that the sine causes firing at the beat frequency of the TI signal it is compared to, but the TI signal needs a higher intensity to fire at its beat frequency.

#### 4.4.2 Steady-state values

From the discussion above, it has become clear that the values of the gates play an important role. The time constants that were discussed play a role in how fast the gates can reach a certain value. The steady-state characteristic then determines to which value the gates evolve. This is investigated in this study by changing the *a*- and *b*-parameters in the logistic function approximation of the gates.

Starting with the m-gate (Figures 6A and 6B), a TI zone is present for all the investigated *a*-values. Increasing the steady-state values shifts the TI zone to higher input currents. For a higher *b*-value, a TI zone appears, while for the *a*-value, the TI zone becomes smaller. The reason for the shift of the threshold to higher input currents is that for higher *a*- or *b*-values, the m-gate is in a more closed state at lower membrane potentials. Interestingly, for values of *b* < 27 mV, the neuron fires spontaneously. In other words, the open probability of the m-gate is high enough at low membrane potentials so the neuron can always enter the positive feedback loop to form an AP.

For the h-gate, the behavior for different values of *a* and *b* can be explained by the fact that for higher *a*-values or lower *b*-values, the h-value around the resting potential is lower. The h-gate is thus already in a more closed state resulting in a lower *Na*^+^-current and a higher input current needed to cause firing.

Useful for the application of TI stimulation is that the steepness of the threshold is not the same for the TI-input and the sinusoidal input. For low *b_h_*-values, this causes a substantial TI zone. The reason for this is the following. When the threshold input is plotted with respect to the beat frequency (Figure 9), it can be seen that a global minimum exists and thus there is an optimal beat frequency for TI stimulation. When the characteristic parameters *a, b* and *k* of the gates are altered, the minimum shifts to other beat frequencies. The beat frequency at the minimal threshold current is shown in Figures 9B-G. For example, increasing the *a_h_*- and *b_h_*-parameter generally shifts the optimal beat frequencies to lower values, with a local maximum at *b_h_* = 10 mV. Hence, the optimal *f_b_* comes closer to 0 Hz and goes further from the case of *f_b_* = 100 Hz with increasing *a_h_*-value. Because the TI signal approximates a sine for *f*_b_ → 0 Hz, a sine can be seen as a TI signal with *f_b_* = 0 Hz, with increasing *a_h_*-value, the threshold for a sine increases less steep than the threshold for the TI signal (Figure 6C). The same holds for the increasing *b_h_* parameter in Figure 6D. In this case, the decrease of the red lines is steeper because the sine comes closer to the optimal beat frequency with increasing *b_h_*. On the other hand, the h-value at low membrane potentials increases which is in turn more favorable to create an AP. This means that there is a double effect for the decreasing threshold. For the 100 Hz beat frequency TI-input, the increase in h-value at low membrane potentials for increasing *b_h_*-value again causes a decrease in threshold but because the optimal beat frequency shifts to lower values, the decrease in threshold is less steep.

**Figure 9.**
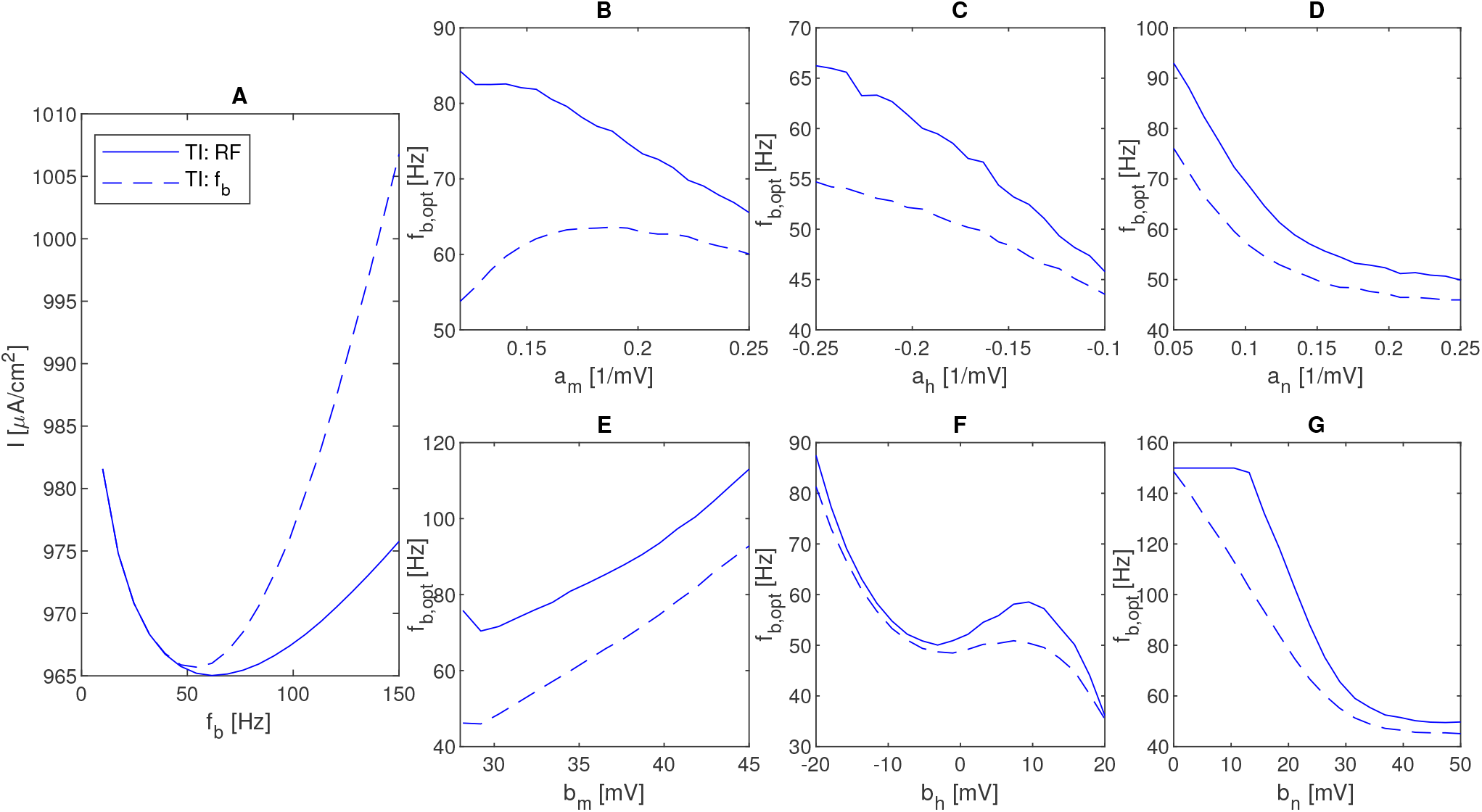
**A**: Illustration of the threshold-beat frequency relation for regular firing (solid line) and firing at the beat frequency (dashed line) for the FH-model with *f_c_* = 3 kHz. **B-G**: Minimal beat frequency for different values of the a and b parameters, where the steady state value of a single gate (x = m, h or n) is substituted by a logistic function. The optimum was found through curve fitting of a fourth order polynomial on the minimum of the simulation and its three closest neighbouring datapoints on each side.

Similarly, the dependency if the optimal *f*_b_ on *a*_X_ and *b*_X_ explains the difference in steepness of the thresholds for the sine and the TI signal for the m- and h-gate. The reasoning can also be followed for the *k*-values. In this case, the effect can be best seen for higher k-values in Figure 5: the optimal beat frequency is inversely proportional with *k*.

The reason for the existence of an optimal beat frequency is because of the following: when the beat frequency is too high, some of the gates might not be restored to a value that is sufficient to create a new AP. Because of this, some beats might be skipped in the neuronal response to the TI waveform. When the beat frequency is too low, the TI waveform envelope builds up too slow. Because of this slow build-up, the h-gate tends to close too fast while the n-gate opens too fast. Because of this the input current needed to create an AP is higher.

Equation (17) can be used to get more insight into the relation between the optimal beat frequency and the membrane channel dynamics. To derive (17), it is assumed that two phases can be distinguished. In each phase *τ_x_* and *x*_∞_ are constant. In the first phase, the gate characteristics take their values at the membrane potential at the peak of the AP. In the second phase, the gate characteristics take their values at the resting potential. This assumption might not be acceptable for the m-gate, which can be approximated as an instantaneous function *m* = *m*_∞_(*V*) due to its fast nature. A second assumption is that when looking at threshold values, the AP is induced at the top of the beat and the gates should be restored around the node of the TI signal. A third assumption is that x does not change much in the half period before the AP. This means these two phases should be ran through in a half period of the envelope 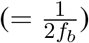. This assumption is supported by our observation in a simplified HH-model, that the h-gate is opened at the envelope node for beat-locked spiking Plovie et al. (2022).

The starting point in the derivation is (2). With the assumption of a constant *τ_x_* and *x*_∞_, the solution for x during the first and second phase is shown in (14) and (15), respectively.

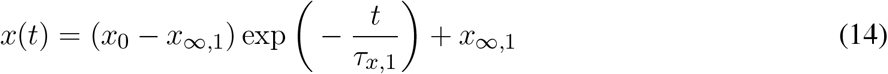

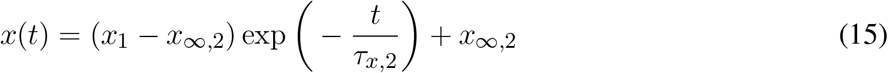

The next step is to solve *x*(*t*_1_) = *x*_1_ for *t*_1_ and *x*(*t*_2_) = *x*_2_ for *t*_2_. The total time is then the sum of *t*_1_ and *t*_2_:

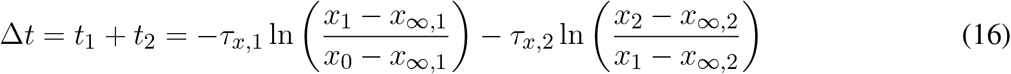

where *x* can be *h* or *n*. Here, *x*_0_, *x*_1_ and *x*_2_ are the gate parameters at the crest, somewhere between crest and trough (at the minimum of the h-gate or the maximum of the n-gate), and at the trough of the envelope, respectively. The optimal beat frequency then becomes:

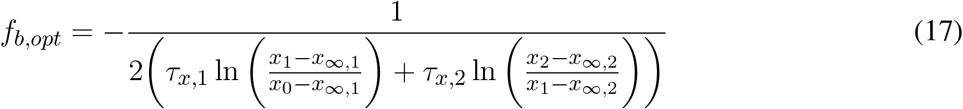

Because *k_x_* is directly related to the time constant (see (10)), a slower gate means that a lower beat frequency is better. This shows the inverse proportionality of the optimal beat frequency with *k_x_*. With this equation, there is an optimal beat frequency for every gate but of course there is only one optimal beat frequency for the whole system. This should be a combination of the different optima for the different gates. Still, (17) gives some mathematical insight into why there is an optimal beat frequency and how it depends on the different gate parameters. As an illustration, when the parameters (*h*_0_ = 0.75, *h*_1_ = 0.025, *h*_2_ = 0.75, *h*_∞,1_ = 0, *h*_∞,2_ = 0.80, *n*_0_ = 0.05, *n*_1_ = 0.56, *n*_2_ = 0.05, *n*_∞,1_ = 1 and *n*_∞,2_ = 0.025) are filled in (17) in the case of spiking at the beat frequency, the resulting optimal beat frequency is 51.1 Hz for the h-gate and 119.3 Hz for the n-gate. From the simulations it was found to be 52.7 Hz for the whole system. When *k_n_* = 3, the optimal beat frequency for the n-gate according to (17) changes to 39.8 Hz while the optimal beat frequency from the simulations is 45.8 Hz. For *k_h_* = 3, the optimal beat frequency for the h-gate according to (17) changes to 17.0 Hz while the simulation gives 23.2 Hz. The optimum is thus inbetween the optimal values of the n-gate and h-gate. The existence of the optimal beat frequency is in line with the concept of resonance in neurons Hutcheon and Yarom (2000). This discussion shows that the resonant beat frequency does not only depend on the time constant of the gates, but also on their steady-state characteristics.

### 4.5 Limitations and future work

Because TI neuromodulation has only recently gained attention, the technique itself can still be improved. For example, the focality of TI stimulation is still in the order of centimeters. This can be improved by adding extra electrode pairs or by using a lensing effect Cao and Grover (2020). Multiple focal points can be achieved by adding extra electrode pairs or by using multiple frequencies in the present electrode pairs. In this way, two TI signals are generated with different carrier frequencies Zhu et al. (2019). Finally, in classic TI stimulation, a reference electrode is placed at the chest. Orientation-tunable temporal interference (ot-TI) is a different method where the reference is chosen locally on the scalp. This heavily influences the electric field lines. With this method, the importance of the direction of the electric field can be shown (Missey et al., 2021).

A first limitation of this study of the underlying mechanism of TI stimulation is that the spatial properties of the electric field are not taken into account in the cell models. Other authors have used morphologically correct neuron models, to investigate the neuronal response to TI modulation Wang et al. (2022). However, axial coupling in these spatially extended models is linear and as a result is not expected to influence the qualitative results obtained in this paper (i.e. dependency of the existence of a TI zone on the GHK nonlinearity and gating dynamics). Nevertheless, morphologically realistic models are interesting to obtain quantitative insights into the activation and conduction block thresholds. Moreover, a limitation of point neurons is that it is not always certain if the detected APs can propagate. A second limitation of this study, is that the TI-modulated point neurons are isolated, while neuronal network simulations have demonstrated the importance of network adaptation Esmaeilpour et al. (2021). As future work, it would be interesting to investigate the influence of synaptic time constants on the TI zone. The underlying mechanism of TI stimulation can thus be found in a lot of aspects of the neural system and may be a combined effect of several factors. Therefore, the investigation in this study of the nonlinearities and time constants in point neurons is a first step in the direction of a comprehensive understanding of neuromodulation by temporal interference.

## 5 CONCLUSION

This study consists of two major parts. First, the discussion of the complexity of neuron models needed to be able to model the experimentally observed responses to TI-DBS. Second, the time dynamics of the ion gates in relation to TI stimulation.

From the first part, it can be concluded that the nonlinearity in the GHK-equation alone is not able to explain the underlying mechanism of TI-DBS. For the FH-model, the linearisation causes minor differences in the neuronal response, that are not sufficient to describe the TI underlying mechanism. For the HH-model, the differences caused by the GHK nonlinearity are bigger. However, the Hodgkin-Huxley TI zone is actually larger in the linearised model, confirming that the GHK nonlinearity is not necessary in order to observe neuronal sensitivity to a beating sine. It should be noted that the choice of the expansion points has a big influence on the neuronal behavior as well. From the investigation of the IF-models it can also be concluded that the introduction of a nonlinear term in the differential equations is not able to shift the threshold of the TI-input beneath that of the sinusoidal input.

From the second part, it can be concluded that the nonlinearities intrinsic to the gating dynamics have a big influence on how the neurons react to the TI and sine input. The mechanism is a complex interplay between the gates that is driven by the gate characteristics in terms of the time constants and steady-state values. This paper showed that changing the gates’ opening and closing rates can influence the existence and the width of the TI zone. A TI zone can appear with increasing *b_m_* values or with a decrease of the *k_m_, a_m_* or *b_h_* parameter. It was also shown that for a certain neuron, it is possible to find the optimal stimulation parameters to increase the TI zone. This is an interesting finding for the optimization of TI stimulation, for which there is a combination of different neuron types at different locations throughout the brain or nerves.

## AUTHOR CONTRIBUTIONS

TP, RS, TT, LM, WJ and ET contributed to the study design. TP performed the simulations and wrote the manuscript with major suggestions of all other authors. RS and TT had a great deal of input into the ideas for simulations. ET supervised the work. All authors read and approved the submitted manuscript.

## ACKNOWLEDGMENTS

We want to thank Michiel De Kesel for the exploratory work and discovering the possible importance of the nonlinearity of the GHK-equation.

