## Supplementary material for "Nonlinearities and Timescales in Temporal Interference Stimulation"

### 1 TAYLOR EXPANSION POINTS

The points for the Taylor expansion for the FH1v2 and gHHv2 models were chosen based on the best average correlation. The resulting values were $0.994$ and $0.971$, respectively. Figure S1 shows the average correlations for all simulated points. The maxima are indicated with a red dot. The location of the expansion point for the gHHv1 model is indicated with a black dot. The location of the expansion point for the FH1v1 model is out of the scope of this plot. It can be seen that a lot of points have a high average correlation and would thus probably not deviate much from the chosen points. On the other hand, there is a region of points with a very bad correlation, especially for the gHH model.

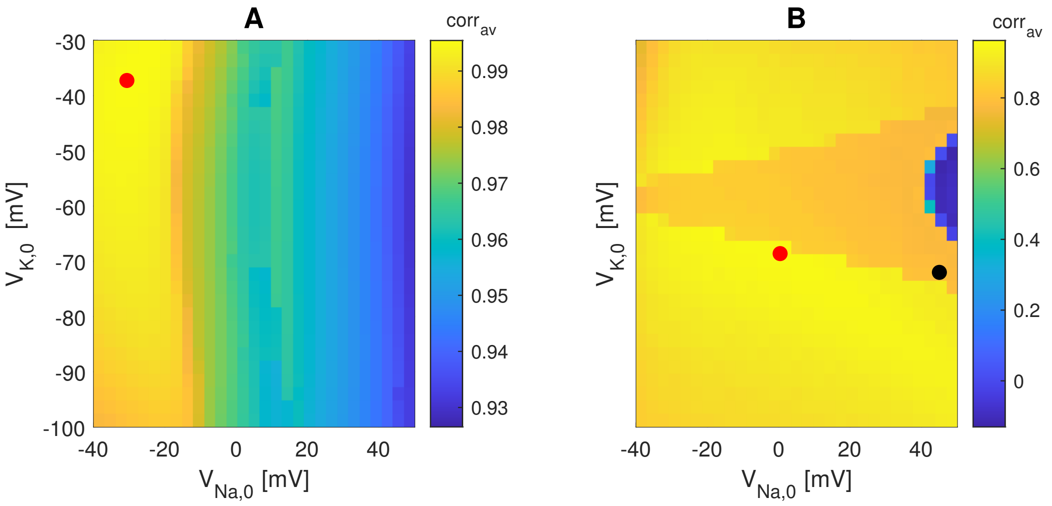


**Figure S1.** Average correlation for the FH1-model and gHH-model compared to the FH- and HH-model for different combinations of the Taylor expansion points $V_{Na,0}$ and $V_{K,0}$. The red dots indicate the maximal correlation. The black dot in the gHH-plot indicates the choice for the gHHv1-model. **A**: FH1-model. **B**: gHH-model.

Some examples of voltage traces with the respective correlations are shown in Figure S2.

### 2 LEAKY INTEGRATE-AND-FIRE NEURONS

It followed from the simulations that leaky integrate-and-fire (LIF) neurons show a small TI zone. This can be explained analytically, starting from the expression of the LIF model:

$$\tau_m \frac{dV}{dt} = RI_e - (V - V_r). \tag{S1}$$

This can be adjusted to:

$$\tau_m \frac{d(V - V_r)}{dt} = RI_e - (V - V_r). \tag{S2}$$

In the frequency domain, this becomes:

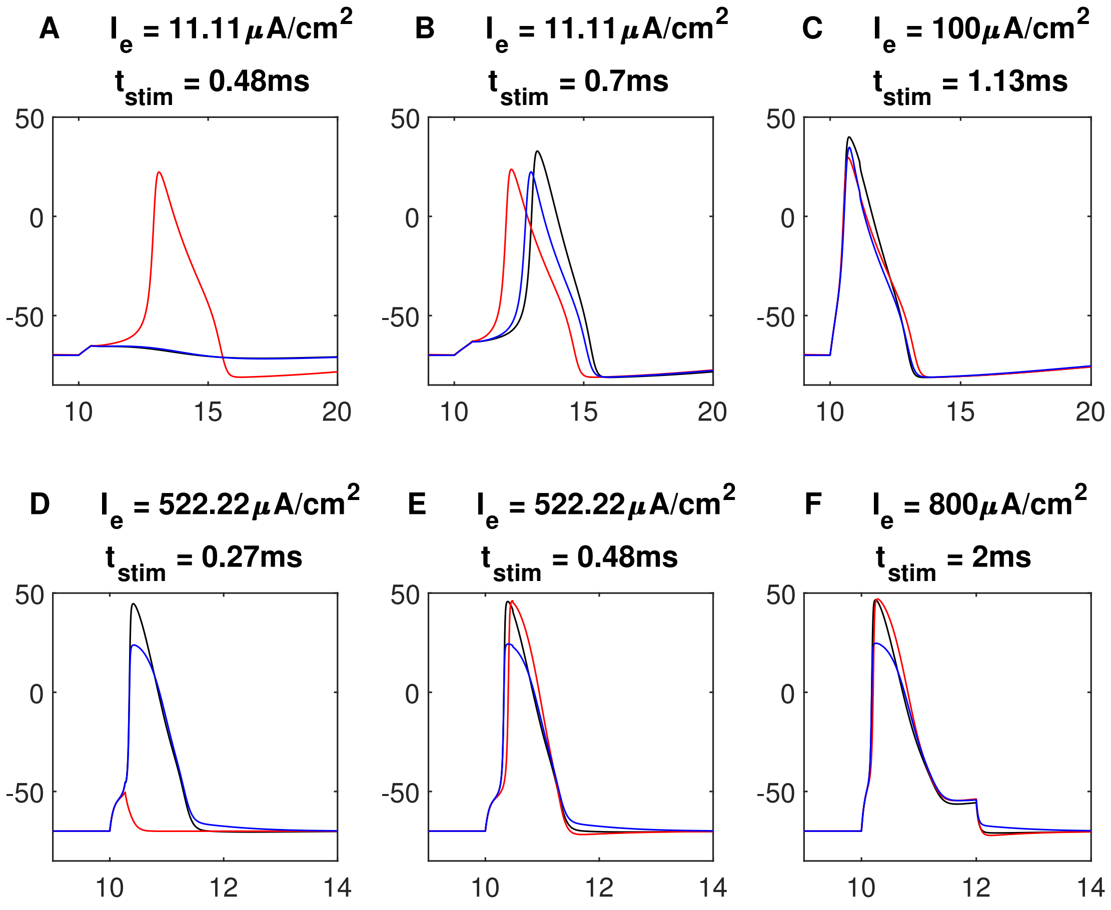


**Figure S2.** Examples of voltage traces used for the calculation of the average correlation between the original Hodgkin-Huxley model and its generalised version (top row) and the Frankenhaeuser-Huxley model and its linearised version (bottom row). The correlations between the black line and the red and blue line is respectively: **A**: 0.449 and 0.996. **B**: 0.638 and 0.942. **C**: 0.994 and 0.998. **D**: 0.248 and 0.994. **E**: 0.955 and 0.994. **F**: 0.992 and 0.994.

$$j\omega\tau_m\tilde{V} = R\tilde{I}_e - \tilde{V}, \tag{S3}$$

with $\tilde{V}$ the Fourier transform of $V - V_r$ and $\tilde{I}_e$ the Fourier transform of $I_e$. The amplitude of $\tilde{V}$ is then

$$|\tilde{V}| = |\frac{R\tilde{I}_e}{1 + j\omega\tau_m}| = \frac{R|\tilde{I}_e|}{\sqrt{1 + (\omega\tau_m)^2}}. \tag{S4}$$

We are interested in the threshold for spiking. $V$ is then $V_{\text{thresh}}$ as defined for the LIF model. Now we have the expression for the threshold of a pure sine:

$$I_{\text{th,ps}} = \frac{V_{\text{thresh}}}{R}\sqrt{1 + (\omega\tau_m)^2}. \tag{S5}$$

The subscript ps stands for pure sine. Because the LIF model is linear, we can apply superposition to find an expression for $|\tilde{V}|$ for the TI signal:

$$|\tilde{V}| = \frac{AR}{2}(|\frac{1}{1+j\omega_1\tau_m}| + |\frac{1}{1+j\omega_2\tau_m}|), \tag{S6}$$

where $A$ is the amplitude of the TI signal. The threshold current then becomes

$$I_{\text{th,TI}} = \frac{2V_{\text{thresh}}}{R}(\frac{1}{\sqrt{1+(\omega_1\tau_m)^2}} + \frac{1}{\sqrt{1+(\omega_2\tau_m)^2}})^{-1}. \tag{S7}$$

The TI threshold is the reciprocal mean of the pure sine thresholds. As a results, the smaller frequency $f_1$ indeed has the bigger influence on the threshold and will drive the threshold to a slightly lower intensity than the pure sine at the carrier frequency.

### 3 EXAMPLE OF A BIG TI ZONE

In the plots of the threshold lines for the FH-model (Figure 3(D, G)), it was seen that the TI zone is very narrow. To apply TI-DBS in reality it would be beneficial to have a wider zone to minimize stimulation in cortical regions. For example on the figure of the threshold as a function of the b-parameter of the h-gate (Figure 6D), it was seen that the width of the TI zone increases with decreasing b-values. As an example, the threshold lines are determined for the case where the steady state of the h-gate is determined by the logistic function with $b_h = -20$ mV and $a_h = -0.18$ 1/mV and the other gates by their original steady state characteristics. The result is shown Figure S3. At $f_c = 3$ kHz, the TI zone has a width of $164.43. \frac{\mu\text{A}}{\text{cm}^2}$. This is 11.97 times bigger than in the normal FH-model. In reality, the technology is of course restricted by the types of neurons that are present in the brain and further research needs to be done to see which neurons are activated as a function of the electric field distribution.

### 4 PHASE DIAGRAMS

Another way to represent the results of a single simulation is with a phase diagram. Here, one of the gates is plotted versus the membrane voltage. Because this model has five variables, this phase diagram is actually a projection on a 2D plane, so it has to be kept in mind that this does not necessarily explain the full behavior of the neuron. Examples are shown in Figure S4. Only the nullclines of the gates (line where $\frac{dx}{dt} = 0$) are shown. Figure S4A shows why the assumption of a constant $m_\infty$ and $\tau_m$ for the derivation of the optimal beat frequency does not hold as the V-range wherein m rises and drops is too wide. For the h-gate and n-gate the oscillations are also over a wide range. However, the oscillations are around $V = -70$ mV which serves as the approximated value.

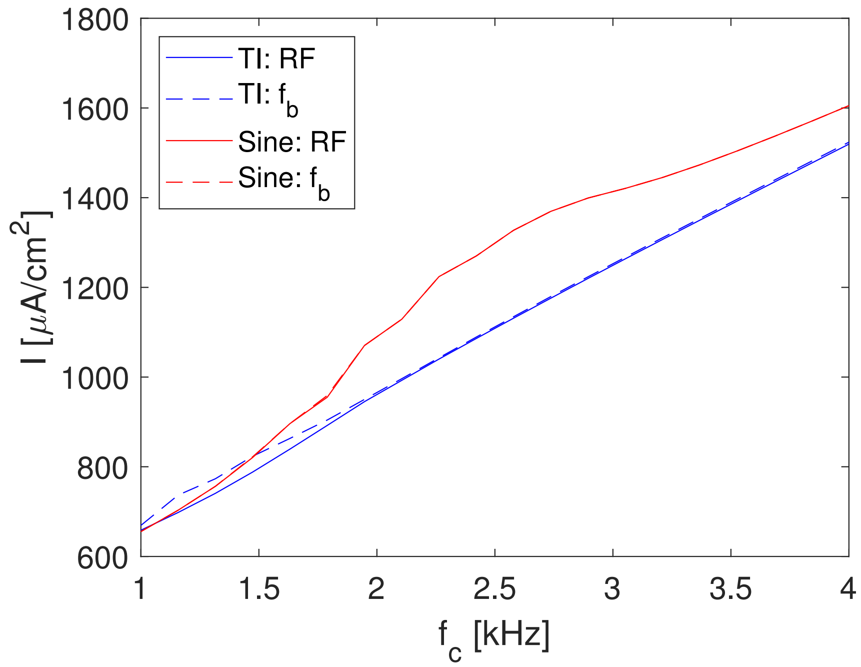


**Figure S3.** Threshold lines of the FH model, adjusted with a logistic function for the steady state characteristic of the h-gate with $a = -0.18\ 1/\mathrm{mV}$ and $b = -20\ \mathrm{mV}$ and a beat frequency $f_b = 100\ \mathrm{Hz}$.

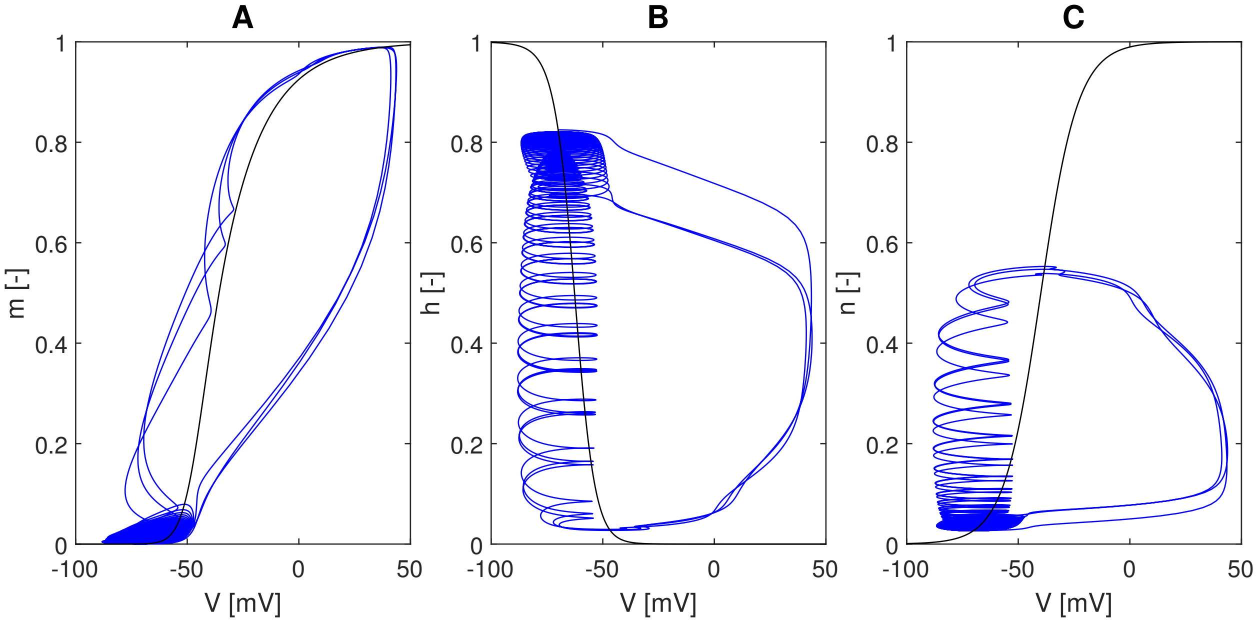


**Figure S4.** Projection of the phase diagram on the $(V, x)$-plane, where $x$ can be $m$, $h$ or $n$. Blue: state of the neuron. Black $x$-nullcline. **A**: m-gate. **B**: h-gate. **C**: n-gate
